# Tumor pre-conditioning of draining lymph node stroma by lactic acid

**DOI:** 10.1101/442137

**Authors:** Angela Riedel, Jonathan Swietlik, David Shorthouse, Lisa Haas, Tim Young, Ana S H Costa, Sarah Davidson, Luisa Pedro, Thordur Oskarsson, Benjamin A Hall, Christian Frezza, Jacqueline Shields

## Abstract

Communication between tumors and the stroma of tumor draining lymph nodes (TDLNs) exists *before* metastasis arises, altering structure and function of the TDLN niche. Transcriptional profiling of fibroblastic reticular cells (FRCs), the dominant stromal population of the LN, revealed reprogramming of these cells in immune related pathways, but also in fibroblast activation and mitochondrial function. However, tumor derived factors driving the changes in FRCs remained to be identified. Taking an unbiased approach, we show that lactate, a metabolite released by cancer cells, elicits upregulation of Pdpn and Thy1 in FRCs of TDLNs, making them akin to activated fibroblasts found at the primary tumor site. Furthermore, we show that tumor-derived lactate alters mitochondrial function of FRCs of TDLNs. Thus, our results demonstrate a novel mechanism by which a tumor-derived metabolite modulates the function of fibroblasts in TDLNs.

## Introduction

The lymphatic system and lymph nodes (LNs) are an integral part of our adaptive immune system and critical for an effective immune response. It is emerging that many tumors exploit lymphatic vessels to spread and colonize downstream LNs, which is an independent indicator of poor prognosis (Cascinelli et al., 2006; Coit et al., 1991; Dadras et al., 2003). The establishment of a pro-tumorigenic environment not only occurs at the primary tumor, but also at sites of metastasis (Joyce and Pollard, 2009). Stromal components of the tumor microenvironment can not only be manipulated by direct cell-cell-interaction with tumor cells, but also by secreted factors. The range of bioactive molecules secreted by tumor cells is diverse, including proteins and peptides (Gelman et al., 2004; Witsch et al., 2010), lipids (Beloribi-Djefaflia et al., 2016; Wang and Dubois, 2010), nucleic acids (Hannafon and Ding, 2013; Schwarzenbach, 2013), exosomes (Peinado et al., 2012), and low molecular weight metabolites such as lactate (Colegio et al., 2014; Warburg et al., 1927; Warburg, 1923). Hence, tumors may use these bioactive factors to prime the metastatic niche environment even before their arrival at the distant site (Peinado et al., 2017). Compared to normal tissue-resident cells, tumor cells exhibit an altered metabolic profile, typically becoming more glycolytic, increasing glucose uptake and lactate secretion (Levine and Puzio-Kuter, 2010). As a consequence, the tumor microenvironment often contains high concentrations of lactate (Walenta et al., 2004). While lactate was originally considered a waste product, its role has been revisited (Gladden, 2004) with the field accepting that most cell types can take up lactate and use it to fuel respiration (Colegio et al., 2014; Faubert et al., 2017; Hui et al., 2017; Yang et al., 2016). An increasing body of evidence shows lactate as a key participant in numerous metabolic pathways, even acting in a signaling capacity (Boussouar and Benahmed, 2004; Chari et al., 2008; Kennedy and Dewhirst, 2010; Philp et al., 2005). In the tumor microenvironment, lactate may act both on tumor cells and supporting stromal cells (Colegio et al., 2014; De Saedeleer et al., 2012; Sonveaux et al., 2012; Vegran et al., 2011). Lactate treatment has been reported to induce hypoxia-inducible factor-1 (Hif-1)-α-driven M2-like polarization of macrophages and concurrent expression of vascular endothelial growth factor (Colegio et al., 2014). Similarly, lactate uptake in normoxic endothelial cells leads to Hif-1α driven angiogenesis (Sonveaux et al., 2012), and cancer-associated fibroblasts (CAFs) can take up and utilize lactate (Rattigan et al., 2012). High lactate levels are also associated with metastasis, tumor recurrence and restricted patient survival in cervical cancers (Walenta et al., 2000). Furthermore, lactate dehydrogenase A (LDHA) expression is upregulated in many cancers and associates with more aggressive tumors. In mouse models, targeting LDHA inhibited tumor growth and lead to delayed metastasis formation (Fantin et al., 2006; Rizwan et al., 2013; Xie et al., 2014). Additionally, LDHA expression was essential for cancer-initiating cell survival (Xie et al., 2014) and has been shown to dampen the tumor immune response of T and NK cells (Brand et al., 2016). Together, these data highlight that metabolic reprogramming does not only impact the tumor cells themselves and metastasis formation, but also the stromal cells in the microenvironment.

We previously demonstrated that fibroblastic reticular cells (FRCs), in TDLNs immediately downstream of murine melanoma undergo remodeling and reprogramming prior to metastatic spread. FRCs are key for LN microarchitecture, but also function as mediators of multiple immunological processes. Transcriptional profiling of FRCs in TDLNs revealed changes in immune related pathways including downregulation of IL7 and CCL21, and in their activation state measured by upregulation of myofibroblast/CAF markers including Pdpn and Thy1. Interestingly, these data also predicted an altered function of FRC mitochondria (Riedel et al., 2016). Despite the observation that FRCs in TDLNs are functionally altered the drivers of changes observed remained undetermined; candidates included mechanical stresses created elevated fluid drainage, tumor-derived cytokines, metabolites or exosomes. Here we propose tumor-derived lactic acid as a driving factor in preconditioning of FRCs of pre-metastatic TDLNs, supporting a change in their metabolic phenotype via alterations to mitochondrial function and induction of myofibroblast-like characteristics. Using computational models, pH was predicted as a key driver of metabolic changes, and a synergistic relationship between lactate and protons as the cause of subsequent fibrotic transformation measured in TDLNs. Tumor derived metabolites indeed decreased intracellular pH leading to increased expression of activation markers Pdpn and Thy1 and impacted mitochondrial behavior of FRCs *in vitro* and *in vivo*.

## Results

### Soluble factors mediate LN expansion and reprogramming

To determine the underlying drivers of the reprogramming of FRCs, we first tested whether the expansion and reprogramming of stroma in TDLNs is mediated by either physical cues, increased drainage, or soluble factors drained from a tumour. To this end we subcutaneously injected PBS, RPMI medium, B16.F10 tumor conditioned medium (TCM) or TCM with and without a 3kDa cut off daily for a period of 11 days into the shoulder of C57BL/6 mice (**Fig 1a**). TCM was sufficient to induce LN swelling and FRC expansion (**Fig 1b**), with soluble factors greater than 3kDa driving this response. Neither PBS nor RPMI media had an impact on the LN or FRCs, indicating that physical cues from increased fluid drainage alone were not responsible for population expansion observed. Pdpn and Thy1 are commonly used as CAF markers and their upregulation in TDLNs is thought to play a role in the pre-metastatic niche establishment. Of note, while complete TCM was able to upregulate Pdpn and Thy1 *in vivo* the impact on both markers was maintained following dialysis (**Fig 1c**), it abolished any effect on FRC number (**Fig 1b**). Though FRC Pdpn protein expression increased in all TCM treatments, Thy1 protein expression was specifically upregulated with low molecular weight TCM (< 3kDa) (**Fig 1c**). *In vitro*, FRCs stimulated with TCM, control conditioned medium from FRCs (CCM), or filtered TCM and CCM with a 3kDa cut-off also showed a significant upregulation of Pdpn and Thy1 at the mRNA level only when treated with TCM and TCM <3kDa (**Fig 1d**) mirroring *in vivo* observations. These data suggested that high molecular weight factors such as secreted growth factors support LN swelling and FRC expansion, whereas small molecular weight factors below 3kDa lead to FRC reprogramming events exemplified by the CAF activation signature described. To corroborate this hypothesis, we performed a cytokine array of tumor secreted factors (**Supp Fig 1a**) and stimulated FRCs with the two most significantly expressed candidates, angiopoeitin-2 and osteopontin. These cytokines had no impact on either Pdpn (**Supp Fig 1b**) or Thy1 (**Supp Fig 1c**) mRNA expression, nor did treatment with TGF-β (**Supp Fig 1d**). Tumor derived exosomes had no impact on these markers (**Sugg Fig 1e**), however there was still a strong induction with exosome depleted supernatant (**Supp Fig 1e**). Further depletion approaches using freeze/thawing, boiling and DNase treatment had no significant effect on Pdpn and Thy1 mRNA levels (**Supp Fig 1f**), indicating that the factors responsible for changes in expression of these markers were small most likely molecule metabolites.

**Figure 1.**
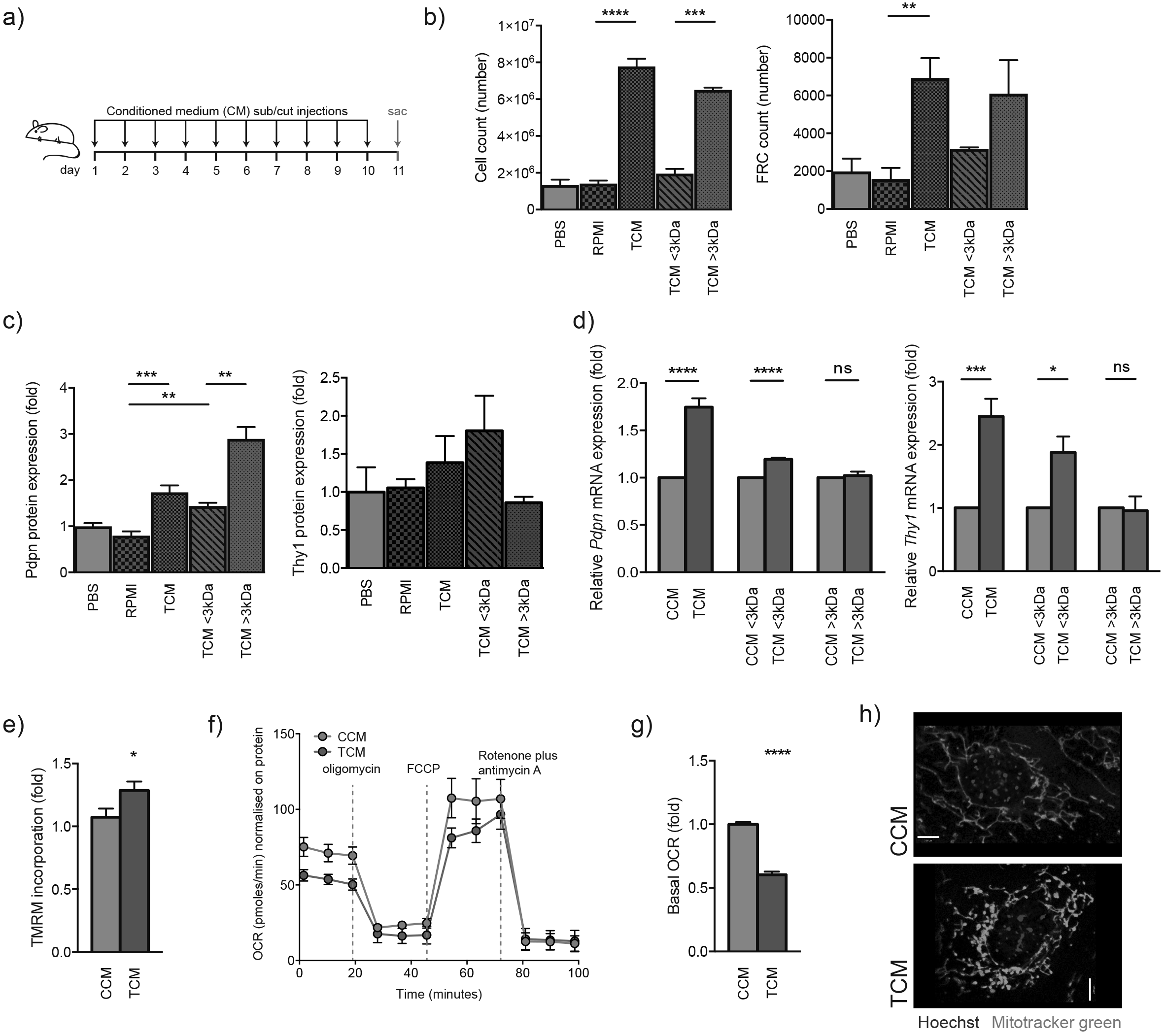
Soluble factors smaller than 3kDa mediate FRC activation and mitochondrial imbalance. (**A**) Experimental scheme used to investigate tumor draining factors *in vivo*. (**B**) Total LN cells (left) or FRCs (right) in PBS, RPMI, tumor conditioned medium (TCM), TCM <3kDa and TCM >3kDa draining LNs from female C57Bl/6 mice assessed by flow cytometry over an injection time of 11 days with sub cutaneous injections each day (mean with SEM +− SD, n = 2-9 animals). (**C**) Pdpn (left) and Thy1 (right) protein expression in FRCs from drainings LNs as in (**A**), assessed by flow cytometry. (**D**) Quantitative RT-PCR analysis of *Pdpn* (left) and *Thy1* (right) in FRCs cultured *in vitro* and treated for 4 days with control conditioned medium (CCM), TCM or with either conditioned medium at <3kDa or >3kDa. (**E**) FRCs treated for 4 days with CCM or TCM, stained with TMRM and analyzed by flow cytometry. (**F**) OCR of FRCs treated with CCM or TCM at baseline and in response to oligomycin, FCCP and rotenone plus antimycin A. One representative experiment. (**G**) Baseline OCR from CCM or TCM treated FRCs. (**H**) Representative confocal images of live cells treated with CCM or TCM. Mitochondria are green (Mitotracker green) and nuclei are blue (Hoechst). *P<0.05, **P<0.01, ***P<0.001 and ****P<0.0001 (two-tailed unpaired *t*-test)

We next sought to determine if TCM exerted a similar impact to FRC mitochondrial function since OXPHOS and mitochondrial dysfunction gene signatures were equally perturbed in TDLN FRCs (Riedel et al., 2016). Indeed, TCM-treated FRCs exhibited increased incorporation of Tetramethylrhodamine, Methyl Ester, Perchlorate (TMRM), a cell-permeant mitochondrial membrane dye that accumulates in active mitochondria, indicating a higher activity of mitochondria in TCM treated FRCs (**Fig 1e**). TMRM is specific for mitochondria with intact membrane potential. Thus to further investigate, we performed a mitochondrial stress test in response to oligomycin, FCCP and a combination of rotenone and antimycin A. In TCM-treated FRCs, the initial oxygen consumption (OCR) was reduced and following addition of FCCP, the maximal respiratory capacity was impaired (**Fig 1f**), and when focusing on basal OCRs, we observed a reduction in TCM-treated FRCs (**Fig 1g**). These changes led us to investigate mitochondrial morphology. Mitotracker green staining and confocal microscopy clearly showed that mitochondria of control (CCM) treated FRCs were elongated and tubular in appearance, whereas mitochondria of TCM-treated FRCs were punctated further indicative of a redox imbalance and altered mitochondrial function (**Fig 1h**).

### Lactate induces mitochondrial and activation reprogramming in LN FRCs

To identify potential tumor-derived factors <3kDa driving the observed functional changes in FRCs, we performed metabolomics analysis of conditioned media using liquid chromatography-mass spectrometry (LC-MS) (**Fig 2a**). While additional metabolites may contribute to TCM composition, lactate and pyruvate dominated as the most enriched metabolites consistent with a glycolytic B16.F10 phenotype. But it was lactate, with more than 10-fold higher levels than any other metabolite which we focused on, confirming the increased levels of lactate observed using enzymatic based assays (**Fig 2b**). Within tumors, Ldha, the enzyme that converts pyruvate to lactate, was detected in close proximity to lymphatic vessels, supporting the hypothesis that tumor derived lactate is drained to TDLNs where it may influence FRC behaviour (**Fig 2c**). Performing further metabolomics analysis using LC-MS of *in vivo* tumor draining lymph nodes (TDLNs), we observed increasing trends in lactate levels in comparison to non-draining lymph nodes (NDLNs), although not statitistically signifncant (**Fig 2d**). Similar trends were also observed in the metabolites succinate and glutamate and of several amino acids (**Fig 2e**). To account for the difficulties in capturing metabolites and metabolite fluxes drained from a tumour at distant sites *in vivo*, we also performed the reverse experiment. To determine whether it was indeed lactate within TCM that was capable of inducing changes in FRC activation or mitochondrial status, we injected 15mM lactic acid (LA) subcutaneously every day for 11 days instead of tumour cells, or TCM *in vivo*. Taking this approach we confirmed that draining lymph nodes reacted to lactic acid by an increase in size (**Fig 2f**) and by the upregulation of *Pdpn* and *Thy1* on mRNA level consistent with TCM reponses (**Fig 2g** and h). Moreover, *in vitro*, FRCs exposed to 15mM LA also displayed significant upregulation of both *Pdpn* and *Thy1* mRNA levels following treatment (**Fig 2i** and j), whereas low pH medium (RPMI pH6) or 15mM sodium lactate, a non acidifying version of lactate had little effect (**Supp Fig 1g** and h). Pdpn upregulation was also verified at the protein level with 15mM LA treatment (**Supp Fig 1i**). As with TCM, LA treatment led to increased incorporation of TMRM (**Fig 2k**) into FRC mitochondria, and mitochondrial stress tests (**Fig 2l**) revealed functional differences, especially in the basal OCR (**Fig 2m**), whereas sodium lactate (SL) and low pH medium did not (**Supp Fig 1j** and k). Moreover, a switch in mitochondria phenotype consistent with those of TCM treatment were detected following LA exposure; elongated mitochondria with vehicle treatment vs. induction of punctated mitochondria with lactic acid treatment (Fig 2n). Interestingly, the mitochondria not only phenotypically changed with LA treatment, but there was also an increase in MTG incorporation measured (**Supp Fig 1l**). Since lactic acid and TCM induced an oxidative and energetic stress, we also performed a viability assay (**Supp Fig 1m**) but no differences were observed consistent with previous studies (Zelenka et al., 2015). In summary these data indicate that tumor-derived lactic acid is sufficient to recapitulate mitochondrial changes and activation signatures induced by TCM in FRCs of TDLNs.

**Figure 2.**
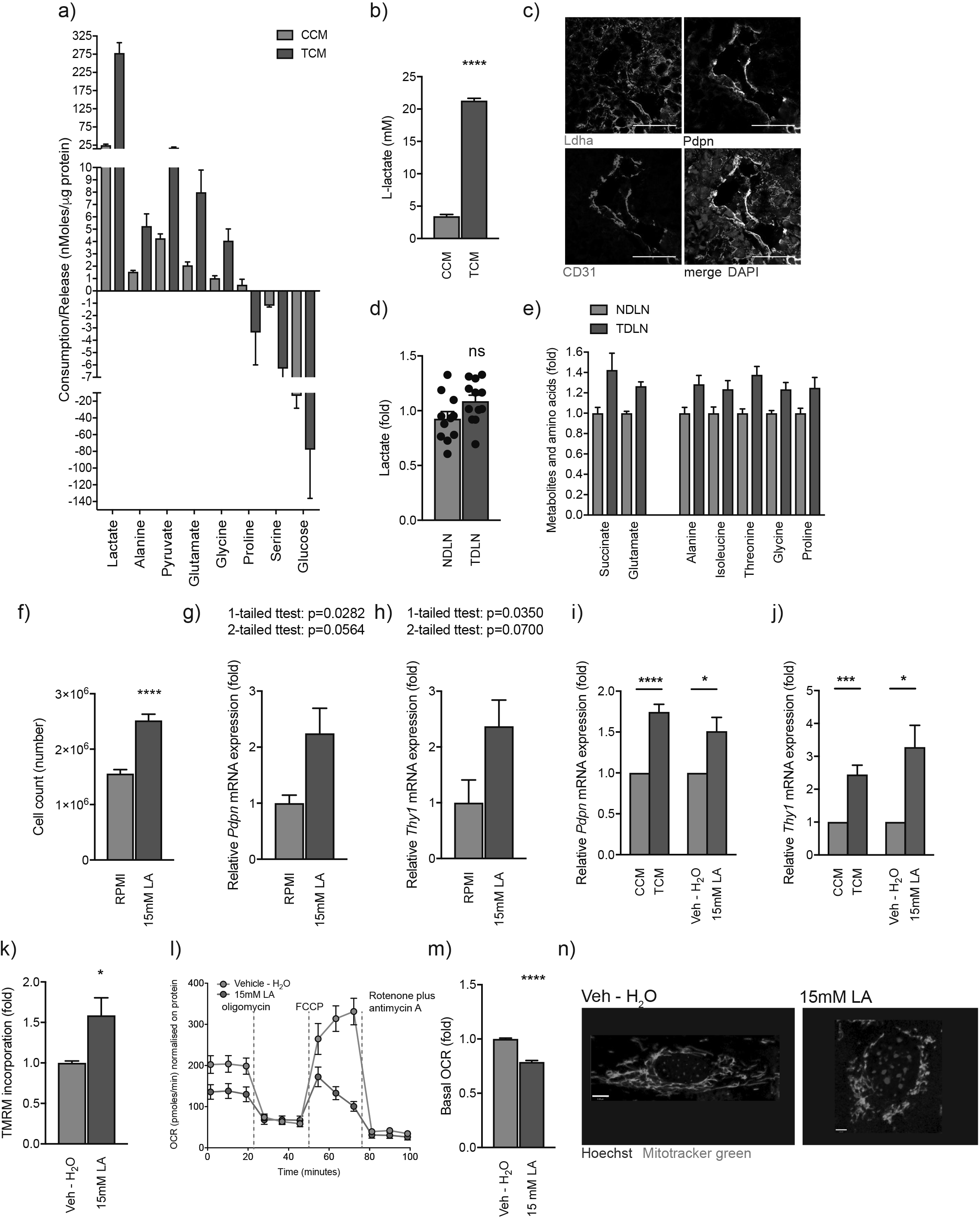
Tumor derived lactate is responsible for observed changes in FRC activation and mitochondria. (**A**) Metabolomic analysis of amino acids and glycolysis- and TCA cycle-intermediates in RPMI supernatant cultured for 24h on FRCs (CCM) or on B16.F10 (TCM) using liquid chromatography mass spectrometry (LC-MS). Displayed are all metabolites with a significant (P<0.05) fold change (FC) ordered according to nMoles/μg protein of cultured cells. (**B**) L-lactate concentration in CCM or TCM measured by an enzymatic assay. (**C**) Representative confocal image of a B16.F10 tumor region. Lymphatic vessel is Pdpn (white) and CD31 (red) positive. Tumor cells surrounding the lymphatics are Ldha (green) positive. Nuclei are in blue. (**D**) Metabolomic analysis of amino acids and glycolysis-and TCA cycle-intermediates in non-draining (ND) and tumour draining lymph nodes (TDLNs) using liquid chromatography mass spectrometry (LC-MS). (**E**) Metabolomic analysis of lactate in non-draining (ND) and tumour draining lymph nodes (TDLNs) using liquid chromatography mass spectrometry (LC-MS). Displayed are all metabolites with a significant (P<0.05) fold change (FC) ordered. (**F**) Total LN cells of RPMI or 15mM LA draining LNs from female C57Bl/6 mice assessed by flow cytometry over an injection time of 11 days with subcutaneous injections each day (mean with SEM +− SD, n = 4 animals). (**G** and **H**) Quantitative RT-PCR analysis of *Pdpn* and *Thy1* mRNA expression in FRCs from draining LNs as in (**F**). FRCs were sorted by flow cytometry from LNs based on their surface expression of Pdpn^+^CD31^−^CD45^−^ (mean with SEM +− SD, n = 4 animals). (**I** and **J**) Quantitative RT-PCR analysis of *Pdpn* and *Thy1* in FRCs cultured *in vitro* and treated with CCM, TCM, Vehicle (Veh – H2O) or 15mM lactic acid (LA) for 4 days. (**K**) *In vitro* FRCs treated for 4 days as in (**I**), stained with TMRM and analyzed by flow cytometry. (**L**) OCR of FRCs treated with vehicle (Veh – H2O) or 15mM LA at baseline and in response to oligomycin, FCCP and rotenone plus antimycin A. One representative experiment. (**M**) Baseline OCR from vehicle (Veh – H2O) and 15mM LA treated FRCs. (**N**) Representative confocal images of live cells treated with vehicle (Veh – H2O) or 15mM LA. Mitochondria are green (Mitotracker green) and nuclei are blue (Hoechst). *P<0.05, **P<0.01, ***P<0.001 and ****P<0.0001 (two-tailed unpaired *t*-test)

### LN FRC activation and mitochondrial changes are Hif1 α independent, but dependent on intracellular pH

To study potential mechanisms of action, we focused on the hypoxia activated transcription factor Hif1α. Tumor-derived lactic acid has been reported to activate Hif1α in both macrophages (Colegio et al., 2014) and endothelial cells (Sonveaux et al., 2012). However, in FRCs, although responsive to hypoxia (1% O_2_), lactic acid (15mM LA) had no impact on Hif1*α* relocalization to the nucleus from the cytoplasm compared with vehicle controls (**Fig 3a**). Moreover, the mRNA levels of Hif1*α* target genes, *Ldha* and *Vegfa* did not change in response to lactate and a significant downregulation of *Gapdh* was observed (**Fig 3b**). Together these results excluded the requirement of a Hif1a-dependent mechanism for lactate-driven FRC reprogramming in TDLNs.

**Figure 3.**
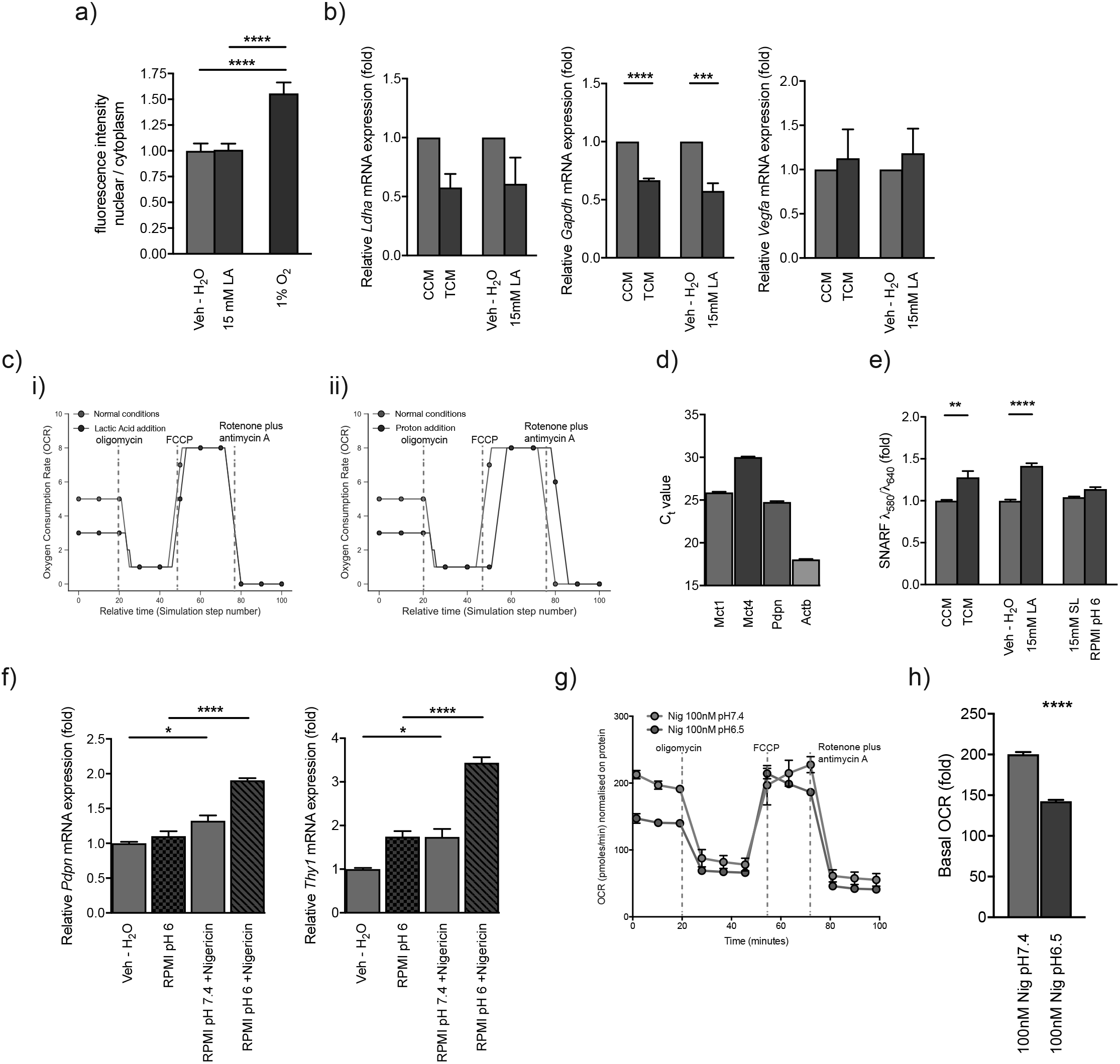
Intracellular pH shift of FRCs contributes to mitochondrial changes and fibrotic signature. (**A**) Quantification of confocal fluorescent images stained for Hif1a and nuclei. Fluorescent intensity was calculated as nuclear / cytoplasm and was normalized on cell number. (**B**) Quantitative RT-PCR analysis of *Ldha, Gapdh* and *Vegfa* in in vitro cultured FRCs treated for 4 days with CCM, TCM, vehicle (Veh – H2O) or 15mM LA. (**C**) Metabolic analysis of mitochondrial function in the computational model. Shown is OCR for mitochondria exposed to lactic acid vs control (i), and mitochondria exposed to low pH vs control (ii). (**D**) Quantitative RT-PCR analysis of *Mct1*, *Mct4*, *Pdpn* and *β-actin* (*Act*) in *in vitro* cultured FRCs displayed in threshold cycle (C_t_) values. (**E**) Intracellular pH of CCM, TCM, Vehicle (Veh – H2O), 15mM lactic acid (LA), 15mM sodium lactate (SL) or RPMI at a pH of 6 (RPMI pH 6) treated FRCs for 4 days and stained with a cell-permeant ratiometric fluorescent pH indicator (SNARF-5F5-(and-6)-carboxylic acid, acetoxymethyl ester, acetate (SNARF)). The indicator exhibits a pH dependent emission shift, which was calculated by *λ*586/*λ*610. Lower intracellular pHs give higher values. (**F**) Quantitative RT-PCR analysis of *Pdpn* (left) and *Thy1* (right) in FRCs cultured in vitro and treated for 48h with vehicle (Veh-H2O, RPMI at normal pH), RPMI at pH 6, RPMI at pH 7.4 with 100nM nigericin or RPMI at pH 6 with 100nM nigericin. (**G**) OCR of FRCs treated with RPMI at pH 7.4 with 100nM nigericin or RPMI at pH 6.5 with 100nM nigericin at baseline and in response to oligomycin, FCCP and rotenone plus antimycin A. One representative experiment. (**H**) Baseline OCR from RPMI at pH 7.4 with 100nM nigericin or RPMI at pH 6.5 with 100nM nigericin treated FRCs. *P<0.05, **P<0.01, ***P<0.001 and ****P<0.0001 (two-tailed unpaired *t*-test)

As tumor-derived LA in lymph nodes impacts several functional pathways in FRCs, it is likely that rather than relying on activation of a single transcription factor, LA induces a more global effect in exposed FRCs. To explore this, we developed a computational model of FRC metabolism under the influence of LA, using an executable model developed from the literature in the BioModelAnalyzer (BMA, **Supp Fig 2a**). We sought to examine the import of lactate and protons, and their corresponding effects on glycolysis and the TCA cycle in response to a simulated metabolic stress test. To simulate a metabolic stress test, nodes within the model, representing either protein activity or membrane properties, were inhibited in sequence, in approximate timings as in the experiment. ATP synthase was inhibited to simulate Oligomycin addition, membrane permeability to protons was altered to simulate FCCP addition, and a node representing mitochondrial complex I through IV was inhibited to represent the addition of rotenone and antimycin A. This gave us the ability to run mitochondrial stress tests *in silico*, and examine the effects of disruption to metabolic pathways on the result. Simulating the addition of LA to the system, the model supported experimental findings that LA reduced the basal OCR significantly in FRCs (**Fig 3c**). We then investigated whether lactate is in fact taken up by LN FRCs. To this end we measured Mct1 (uptake) and Mct4 (secretion) mRNA expression levels on FRCs *in vitro*. Interestingly, Mct1 exhibited higher levels of expression than Mct4 (**Fig 3d**). As Mct1 is a proton-coupled monocarboxylate transporter, lactate must be taken up from the extracellular space together with a proton, which leads to a decreased intracellular pH. Thus, intracellular pH of FRCs was measured using the cell-permeant ratiometric fluorescent pH indicator (SNARF-5F5-(and-6)-carboxylic acid, acetoxymethyl ester, acetate (SNARF)). TCM and 15mM LA(15mM LA) treatment of FRCs induced a significant decrease in pH compared with CCM, 15mM sodium lactate (15mM SL) or pH 6 medium (RPMI pH 6) evidenced by an increased SNARF emission shift (**Fig 3e**). Upon further interrogation, the model predicted that addition of protons alone was enough to replicate the effects of LA on mitochondria, whereas sodium lactate addition was not enough to alter the OCR (**Supp Fig 2b**). To further validate the model’s predictions, we examined potential changes induced by a sudden decrease of intracellular pH in LA treated FRCs *in vitro*. We mimicked the LA scenario with the help of the potassium ionophore nigericin (Nig), which is a K^+^/H^+^ exchanger and commonly used to adjust intracellular to extracellular pH. As nigericin, a microbial toxin, has been shown to induce apoptosis via Caspase-1 (Hentze et al., 2003) and reported to disturb the cell’s mitochondrial bioenergetics by matrix acidification leading to a reduced basal respiration (Manago et al., 2015), we used very low concentrations and short treatment durations of this drug in the following combinations: normal pH media, low pH media, normal pH media plus nigericin or low pH media plus nigericin. We detected an increase in mRNA expression levels of *Pdpn* and Thy1 with nigericin and normal pH medium alone. This was further augmented with nigericin and a low pH medium, indicating that the induction of *Pdpn* and *Thy1* is dependent on the intracellular pH and associated mitochondrial functional changes (**Fig 3d**). Intracellular pH was critical for proper mitochondrial function since FRCs treated with nigericin and a low pH medium presented with a decrease in the basal OCR (**Fig 3e** and **f**). Adjusting intracellular pH alone mirrored the situation of lactic acid uptake, impacting intracellular pH and both mRNA expression levels of *Pdpn* and *Thy1* and mitochondrial function.

### Lactic acid induces LN FRC fibrotic signature

To identify additional relevant deregulated components in LN FRCs treated with TCM and LA, we performed a PCR array specific to fibroblast function/pathology (Mouse Fibrosis, QIAGEN). Of the 84 mRNA transcripts that were measured, TCM treatment significantly changed 26%, lactic acid 27% and filtered TCM (<3kDa) 29% of them, with a significant proportion overlapping between the 3 different treatments (**Fig 4a**). This was especially true for all upregulated mRNAs (**Fig 4b**), revealing large similarities between lactic acid and TCM treatments. Among the deregulated mRNA transcripts were growth factors including *Agt, Hgf* and *Tgfb2*, ECM components and remodeling enzymes *Dcn, Col1a2* and *Mmp2* (**Fig 4c**). mRNA levels of the transcription factor Stat1 and the integrin Itga1 were also upregulated. Acta2 (α-smooth muscle actin), another marker for activated fibroblasts (Kalluri, 2016) was also significantly upregulated with filtered TCM (<3kDa) showing the same tendency for TCM and 15mM LA. Among the downregulated mRNA transcripts were *Gapdh, Timp1* and *Itgb5* (all **Fig 4c**). Although *Col1a2* mRNA was significantly upregulated by 15mM LA, but not with TCM treatment, confocal imaging confirmed a significant increase in collagen1 at the protein level in FRCs treated with both 15mM LA, and TCM treatment (**Fig 4d**).

**Figure 4.**
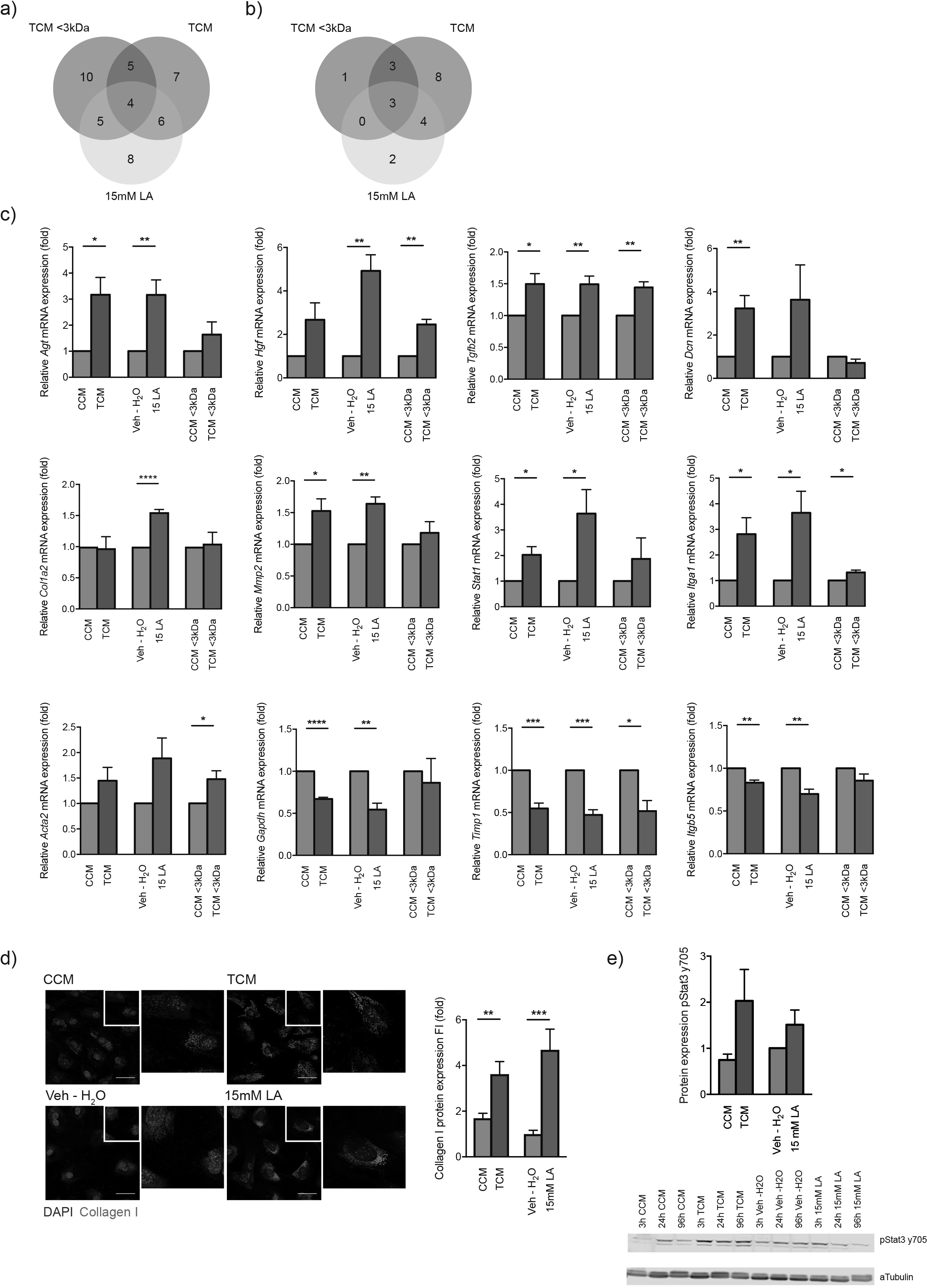
Tumor factors and lactic acid induce LN fibrosis. (**A**) Overlap of genes covered by the PCR array (PAMM-120ZG, Mouse Fibrosis, QIAGEN) significantly (P<0.05) deregulated in comparisons between TCM to CCM, TCM <3kDa to CCM <3kDa and 15mM LA to vehicle (Veh – H2O). Each sample was run 3 times from 3 independent biological replicates. (**B**) Overlap of genes significantly (P<0.05) upregulated in comparisons as in (**A**). (**C**) Quantitative RT-PCR analysis (PCR array data) of *Agt, Hgf, Tgfb2, Dcn, Col1a2, Mmp2, Statl, Itgal, Acta2, Gapdh, Timp1* and *Itgb5*. (**D**) Confocal images stained for collagen I (red) and nuclei (blue; left) and quantification (right). Scale bar = 50μM. (**E**) Western blot (bottom) for pStat3 y705 of lysates from 3h, 24h and 96h treated in vitro cultured FRCs with CCM, TCM, vehicle (Veh – H_2_O) or 15mM LA and quantification of the 3h time-point (top). αTubulin as loading control. *P<0.05, **P<0.01, ***P<0.001 and ****P<0.0001 (two-tailed unpaired *t*-test)

Signal transducer and activator of transcription (Stat) 3 has been implicated in fibrotic conditions (Chakraborty et al., 2017) and is reported to play a role in converting fibroblasts to proinvasive cancer-associated fibroblasts (Albrengues et al., 2015). Since Pdpn has a Stat3 binding site in its promoter regions (Honma et al., 2012), and lactate has been shown to lead to Stat3 and ERK phosphorylation in human mesenchymal stem cells (Rattigan et al., 2012) we next sought to determine Stat3 signaling levels in FRCs in response to lactic acid exposure. An upregulation of pStat3 y705 was observed in early time points (3h) in both TCM and LA treated samples, compared to CCM or vehicle treated samples, indicating LA-induced activation of Stat3 signaling in these cells, upstream of Pdpn induction (**Fig 4e**).

Where possible, we incorporated nodes into our model representing the altered expression levels of the fibrotic/activation signature. The model also supported *in vitro* experiments confirming LA mediated transformation of FRCs towards a fibrotic phenotype. In particular, Tgfb2 mediated activation of Smad can explain the transformation, leading to activation of Mmp’s, collagen synthesis, and Pdpn expression. In summary, the model implied that lactate and a dissociated proton are co-transported into the cell (**Supp Fig 2c**), a build-up of intracellular lactate inhibits the activity of lactate dehydrogenase (LDH), and the drop in intracellular pH inhibits phosphofructokinase (PFK). Secondly, protons are transported via ATP synthase into the mitochondria, where the resultant reduction in pH inhibits the activity of enzymes involved in the TCA cycle. Additionally, intracellular lactate and protons synergistically activate Tgfβ2, and the transcription factors Smad and Stat3, among others, which upregulate Pdpn, collagen synthesis, Timps, and Mmps. Major enzymes are represented as blue boxes, and disruptions to normal activity are represented as black arrows.

## Discussion

Lymph node metastasis is a major cause of cancer-related mortality in many tumors. To accommodate and support development of secondary tumors, lymph nodes adapt, establishing a pro-tumor pre-metastatic niche. We previously demonstrated that lymph nodes immediately downstream of tumors enlarge, and stromal populations remodel and undergo transcriptional reprogramming in response to tumor-derived cues ***prior*** to metastasis (Riedel et al., 2016), yet the cues responsible for these adaptations remained unclear. Here we show that instead of biophysical cues such as increased flow or large proteins and nucleic acids, it is the tumor-derived metabolite lactate that induces transcriptional reprogramming of two pathways key to FRC function: activation status and mitochondrial dysfunction.

Acidification and lactate secretion in the tumor microenvironment (TME) is associated with cancer progression and metastasis (Brizel et al., 2001; Walenta et al., 2000), and potentially acquisition of therapeutic resistance (McCarty and Whitaker, 2010). Thus, increasing the extracellular pH of primary tumors could have therapeutic impact. However, a direct link between lactate, pH and the establishment of a pre-metastatic niche to support metastases has not yet been identified. Our data indicates that lactate secreted by tumor cells impacts not only the local tumor, but also the downstream draining lymph node. Lactate and proton uptake by FRCs in the LN led to acquisition of a pro-fibrotic, activation signature and altered mitochondrial structure coupled to reduced function likely mediated by a change in intracellular pH. Both fibrosis and changed metabolism of stromal cells in the TME have been connected to tumor progression and metastasis (Cox and Erler, 2014; Kalluri, 2016; Xing et al., 2015). Within the local TME, lactate has been reported to induce a M2-like phenotype in stromal macrophages, accompanied by VEGF expression and angiogenesis and driven by Hif-1α activation (Colegio et al., 2014; Sonveaux et al., 2012). Sonveaux and colleagues demonstrated that Hif1α depended on pyruvate levels and Phd activity, requiring 2-oxoglutarate as a substrate; increased levels of pyruvate following uptake and conversion of lactate to pyruvate by Ldh led to Phd inhibition and Hif1α protein stabilization (Sonveaux et al., 2012). In contrast, in our hands, the observed effects on FRCs of TDLNs were not Hif-1α dependent. This might be a result of several factors; Colegio et al. used much higher concentrations of lactic acid, up to 40mM compared with 15-20mM measured and used in the present study (Colegio et al., 2014), and Sonveaux et al. used sodium salts to buffer all media at pH 7.3, hence any pH effect of lactate would not be apparent (Sonveaux et al., 2012). Moreover, Hif-1α dependency is likely to be context and cell-type specific. Consistent with our previous study where immune populations in TDLNs were altered with T cell survival effected, Brand et al. showed that lactic acid accumulation in the TME led to impaired function and survival of T and NK cells via intracellular acidification, contributing to the tumor immune escape. Furthermore, they correlated LDHA mRNA expression with worse overall survival in a cohort of 44 metastatic melanomas (Brand et al., 2016).

In the present study we observed the induction of an activated, fibrotic phenotype in TCM and lactic acid treated FRCs both *in vitro* and *in vivo* consistent with work by Kottmann et al., which showed a role for lactic acid in myofibroblast differentiation in idiopathic pulmonary fibrosis (IPF). Here, lactic acid accumulation and LDH5 expression in IPF lungs activated TGF-β via a pH-dependent mechanism to induce myofibroblast differentiation, αSMA and Calponin expression coincident with collagen and other ECM protein deposition (Kottmann et al., 2012). Previously, we observed that FRCs of TDLNs become more activated reminiscent of cancer associated fibroblasts (CAFs) and fibroblasts in fibrotic disease. This was evident by not only increases in Pdpn and Thy1 expression, but also by an altered production of collagen and an increased contractility. Measured in high quantities in TCM and within tumours *in vivo*, we identified lactic acid as a driver of Pdpn and Thy1 expression and collagen production by FRCs in downstream draining LNs. Although direct sampling and quantification of lactate within TDLNs *in vivo* was variable, the trends observed warranted further investigation. Indeed, a peripheral subcutaneous bolus of lactate alone was sufficient to induce the same FRC activation signatures in draining nodes as measured in TDLNs (Riedel et al., 2016), *in vitro* and with TCM doses *in vivo*. Consistent with this, lactate has been shown to induce collagen genesis in fibroblasts (Green and Goldberg, 1964). Furthermore, the most deregulated mRNA transcripts by LA presented here including *Agt, Hgf, Mmp2* and *Tgf-β2*, are known to be deregulated in pathological fibrosis (Kendall and Feghali-Bostwick, 2014) and pro-tumorigenic cancer associated fibroblasts (Kalluri, 2016; Kalluri and Zeisberg, 2006).

In addition to changes in the activation state of FRCs, we observed mitochondrial adaptations in TCM and LA treated FRCs. Zhang et al. recently showed that CAFs induced with either TGF-β1 or PDGF have reduced oxidative phosphorylation, which was IDH3α dependent (Zhang et al., 2015). Here, a HIF1α dependent metabolic switch from oxidative phosphorylation to aerobic glycolysis was induced. In concordance with our results, basal oxygen consumption rates in newly converted CAFs were significantly decreased (Zhang et al., 2015). Moreover, Buck et al linked mitochondrial dynamics with T cell fate where fissed mitochondria (punctate) exhibited imbalanced redox, and fused mitochondria (elongated, tubular) mitochondrial function (Buck et al., 2016); T_M_ cells possessed fused mitochondria and T_E_ cells fissed mitochondria. As for our data in FRCs, the punctate mitochondrial phenotype of T_E_ cells coincided with increased incorporation of MTG and a decreased OCR and spare respiratory capacity (SRC). However, no link has been established so far between low intracellular pH, lactate and a changed mitochondrial phenotype in T cells.

To summarize, the present study set out to decipher the influence of the primary tumor on draining LN and the stromal cells within, defining tumor-derived factors responsible for pre-metastasic adaptations. From this study it is clear that not one single factor is responsible for all tumor induced FRC effects such as proliferation and immune signatures: both areas that warrant further investigation. However, while excluding mechanical cues such as elevated fluid drainage, or proteins and nucleic acid, we identified lactic acid as one of the major factors inducing stromal cell reprogramming towards more activated and metabolically altered statuses. The functional changes induced in stromal cells highlights the potential benefit of targeting tumor acidity and/or lactate accumulation both in the primary tumor itself but also the extended tumor microenvironment, the draining lymph node.

## Author contribution

A.R. designed, planned and performed most experiments and associated analyses. J.S. and L.H. helped with the *in vitro* experiments. L.P. and S.D. participated in western blot analysis and tissue stainings. S.C., T.Y. and C.F. advised on, conducted and analyzed data of the metabolomics studies. D.S. contributed with a model for FRC metabolism. C.F. and B.A.H. provided intellectual feedback and support. A.R. and J.S. conceived of the project, guided research and interpreted data. A.R., D.S. and J.S. wrote the paper. All authors revised the manuscript.

## Acknowledgements

We thank the members of the Shields and Frezza Group for comments and discussion; members of the Ares Facility E23 staff for animal husbandry and technical support;

Supported by Medical Research Council core funding (J.S. and C.F.) and the Royal Society (UF130039 to B.A.H.).

## Experimental procedures

### Animal experiments

All experiments involving mice were performed in accordance with UK Home Office regulations under Project License PPL 80/2574. All experiments were performed on immune competent 8-9 week old female C57BL/6 mice. Conditioned medium, RPMI, PBS or 15mM lactic acid were injected subcutaneously into the shoulder region for 11 days every day and animals were culled and lymph nodes (LNs) retrieved on day 11. For syngeneic tumors, 2.5 × 10^5^ B16.F10 melanoma cells were inoculated subcutaneously into the shoulder region and animals were culled and tumors retrieved at day 11. Animals were only excluded if tumors failed to form or if health concerns were reported. Tumor size was monitored with calipers and the volume was calculated based on the ellipsoid formula π/6 × (length × width^2^).

### Flow cytometry

Extracted LNs from mice were mechanically disrupted and digested in a 500 μl mixture of 1 mg/ml collagenase A and D (Roche) and 0.4 mg/ml DNase I (Roche) in PBS at 37 °C for 30-60 min with 600 rpm rotation. EDTA (final concentration 10 mM) was then added and cells were passed through a 70 μm mesh prior to immunostaining. *In vitro* cell cultured cells were detached by either Trypsin-EDTA (Gibco, Life Technologies) or Accutase (Sigma). Cells were stained with fixable viability dye live/dead violet (Molecular Probes) and combinations of the following fluorescently conjugated antibodies; Pdpn (8.1.1), Thy1 (30-H12) (both BioLegend). For intracellular staining FoxP3/Transcription Factor Fixation/Permeabilization Kit (eBioscience) guidelines were followed.

For mitochondrial stainings, cells were stained with MitoTracker Green FM (MTG; Cell Signaling) and Tetramethylrhodamine, Methyl Ester, Perchlorate (TMRM; Thermo Fisher Scientific) at the concentration of 50nM (MTG) and 20nM (TMRM) in treatment/conditioned medium for 30 min at 37°C and 5% CO_2_.

Intracellular pH was measured with the help of a cell-permeant ratiometric fluorescent pH indicator called SNARF (SNARF-5F5-(and-6)-carboxylic acid, acetoxymethyl ester, acetate; Thermo Fisher Scientific). This SNARF pH indicator is excitable at 532nm and exhibits a pH dependent emission shift from 580nm to 640nm. The cells were stained with 10nM SNARF in treatment/conditioned medium for 30 min at 37°C and 5% CO_2_ and analyzed using a LSR Fortessa with a 532 laser and 586/15 and 620/20 filters. Emission shifts could then be calculated based on λ_586_/λ_610_ using the geometric mean with lower intracellular pHs giving higher values. All flow cytometry was performed on LSR Fortessa (BD Biosciences) analyzers and offline analysis was performed with FlowJo software (Treestar).

### Cell sorting followed by RNA analysis

For RNA processing, LN cell suspensions were sorted on a High speed Influx Cell Sorter (100 μm nozzle, BD Biosciences) into RNA protect Cell Reagent (QIAGEN). RNA was isolated with the RNeasy plus micro Kit (QIAGEN) and RNA quality and quantity was analyzed with a Bioanalyzer (Agilent Technologies). Only RNA samples with a RIN value above 8 and total concentration of 100 pg/μl were further processed for whole transcriptome amplification via the Ovation PicoSL WTA V2 Kit (NuGEN). Quantitative RT-PCR was performed using TaqMan assays (*Pdpn* Mm01348912_g1, *Thy1* Mm00493682_g1, *Actb* Mm00607939_s1) and a StepOne Real Time PCR System instrument (both Life Technologies).

### Cell Culture

B16.F10 (CRL 6475, ATCC) cells were maintained in DMEM with 10 % fetal bovine serum (Sigma Aldrich) and 100 U/ml penicillin-streptomycin (both Life Technologies). A FRC cell line was isolated from p53^ER/ER^ C57BL/6 mice and characterized based on their expression levels of Pdpn and VCAM-1 and their lack of expression of CD45 and CD31. FRCs were then *in vitro* cultured in RPMI-1640 medium supplemented with 100 U/ml penicillin-streptomycin (both Life Technologies), 10 % fetal bovine serum (FBS, Sigma), 10 mM HEPES and 15 μM β-mercaptoethanol (Sigma). All cells were incubated at 37 °C with 5 % CO_2_. For tumor cell conditioned medium (TCM) or control conditioned medium (CCM) production; B16.F10 cells or FRCs were seeded at 20% confluency into 175cm^2^ culture flasks, after 24h the medium was exchanged for RPMI-1640 (Sigma) without supplements. After another 24h the medium was retrieved and filtered (pore size 0,22 μm) and frozen at −80°C for storage. Conditioned media was never subject to repeated freezing/thawing. For treatment studies, 5,000 FRCs were seeded into a 6-well plate in full growth medium, 24h later the medium was exchanged for 100% conditioned medium or medium containing 15mM lactic acid or sodium lactate (both Sigma) supplemented with 2% FBS and 100 U/ml penicillin-streptomycin. After 48h the medium was exchanged for the same conditioned medium and after 96h the cells were harvested for analyses. RNA extraction was performed using RNeasy Mini Kit (QIAGEN). 1 μg of RNA was used for reverse transcription using First Strand cDNA synthesis Kit (Thermo Scientific). Thereafter, qRT-PCR was performed using TaqMan assays (*Pdpn* Mm01348912_g1, *Thy1* Mm00493682_g1, *Mct1* Mm01306379_m1, *Mct4* Mm00446102_m1, *Ldha* Mm01612132_g1, housekeeping gene *Actb* Mm00607939_s1) and a StepOne Real Time PCR System instrument (both Life Technologies).

### Immunofluorescence microscopy

FRC were seeded on 8 chamber glass slides and treated over different periods of time. Chamber and tissue slides were then fixed in either 4% PFA for 10 min at RT or a 1:2 Acetone:Methanol solution for 5 min at −20°C. Cells were permeabilized with 0.2 Triton X-100 (Sigma) in PBS and incubated for 5 min on ice. Followed by a blocking step for 40 min in 5 % chicken serum, 2 % BSA, 0.1% Tween20 in PBS at RT. Finally, the cells were incubated with the following primary anti-mouse antibodies at 4 °C overnight: Collagen I (Abd Serotec), Hif1a (ERP16897, Abcam), Ldha (Abcam), CD31 (MEC13.3, BioLegend), Pdpn (8.1.1, BioLegend) and DAPI nuclear counterstain. For all stainings, Alexa Fluor secondary antibodies (Thermo Scientific) were used and slides were mounted in ProLong Gold (Thermo Scientific). For life cell imaging of mitochondria, FRCs were stained with Hoechst 33342 (Life technologies) and 50 nM MitoTracker Green FM (Cell Signaling) for 30 min at 37°C and 5% CO_2_ and imaged live after a washing step in RPMI medium with the respective treatment. All confocal images were taken using either a Leica SP5 or Zeiss LSM 880 confocal microscope and processed with Volocity (Perkin Elmer) or FIJI (ImageJ).

### Metabolic assays

Oxygen consumption rate (OCR) and extracellular acidification rate (ECAR) were measured using an Extracellular Flux Analyzer, Seahorse XF^e^14, system (Seahorse Bioscience). 30,000 FRCs were seeded into each well of a 24-well plate, incubated at 37°C and 5% CO_2_ overnight and directly before the assay the medium was exchanged by bicarbonate-free RPMI (Sigma) and, where appropriate, 15mM lactic acid or conditioned medium. Cells were then incubated in a CO_2_-free incubator at 37°C for 30 min. OCR and ECAR were determined under basal conditions and in response to 1 μM oligomycin, 1 μM fluoro-carbonyl cyanide phenylhydrazone (FCCP) and 1 μM rotenone + 1 μM antimycin A. Protein content (BCA assay, Thermo Scientific) was used for data normalization.

### PCR array

The RT^2^ Profiler PCR Array PAMM-120ZG (Mouse Fibrosis, QIAGEN) was used. RNA was extracted using a RNeasy Mini Kit (QIAGEN) with an additional DNA digestion step. 400ng of total RNA was then reversely transcribed using the RT^2^ First Strand Kit (QIAGEN), followed by Real-Time PCR using SYBR Green technology and a Roche LightCycler 480. Each gene was normalized on recommended housekeeping genes and fold changes were calculated based on sample replicates.

### Immunoblotting

Cell pellets were lysed in Pierce-RIPA buffer (Thermo Scientific) supplemented with protease-inhibitor cocktail (Roche), 1mM PMSF and 1mM Na_3_VO_4_. Protein concentration was measured by BCA assay (Thermo Scientific), samples were boiled at 99°C for 5 min in protein loading buffer and loaded at 25 μg. Samples were then separated by a 12% SDS-PAGE and transferred onto a Nitrocellulose membrane (Millipore). The membranes were blocked with Odyssey blocking buffer (Li-Cor) for 1 hour and then incubated with a pStat3 (Phospho-Tyr705, Cell Signaling) or α-tubulin (B-5-1-2, Sigma) primary antibodies overnight at 4°C in 5% BSA (w/v) in PBS and 0.1% Tween 20. After washing with PBS, membranes were incubated with the appropriate fluorescent-conjugated secondary antibody (IRDye 680RD Donkey antiMouse – 925-68072; IRDye 800CW Donkey anti-Rabbit – 925-32213) for 1 hour. Detection was performed using Odyssey CLx (Li-Cor).

### Lactate measurements

L-Lactate was measured in conditioned medium with the help of the L-Lactate Assay kit (ab65331, abcam) and were done according to the manufacturer’s instruction. All samples were beforehand deproteinized with the Deproteinizing Sample Preparation Kit – TCA (ab204708, abcam).

### Sulforhodamine B (SRB) colorimetric cell density assay

Cells were fixed in 1% (or 10%) trichloroacetic acid (TCA) (v/v) in H2O for 30 min and stained with 0.057% (w/v) SRB stain (Fluka) in 1% (v/v) acetic acid for 30 min. Thereafter the plates were washed 3 times in 1 % (v/v) acetic acid and dried overnight. Finally, it was resuspended in 10mM Tris base and OD was measured at 510nm in a platereader (Tecan).

#### Exosome isolation/depletion by ultracentrifugation

To remove cells and cellular debris, conditioned medium was centrifuged for 20 min at 2000 × g and 4 °C. The supernatant was recovered and transferred into polyallomer tubes suitable for ultracentrifugation. Following centrifugation for 30 min at 10000 × g, 4 °C, the supernatant was transferred to a fresh tube and centrifuged for 70 min at 100,000 × g, 4 °C. The supernatant represented exosome-free medium and was removed leaving 2 mm of liquid above the pellet. To wash the isolated exosomes, the pellet was resuspended in 1 mL PBS. Then PBS was added to fill the tube completely. After centrifugation for 1 h at 100,000 × g, 4 °C, the supernatant was removed and the pellet was resuspended in PBS and stored at −80 °C.

### Factor depletion

#### Heat denaturation

Conditioned media were incubated at 99 °C for 15 min.

#### Freeze and thaw cycles

Conditioned media were frozen at −80 °C for 15 min and thawed at 60 °C for 15 min. This was repeated three times.

#### Filtering (molecular weight cut-off (MWCO) = 3 kDa)

Conditioned media were filtered at 4000 × g, 4 °C using Amicon Ultra-4 Centrifugal Filter Units (Merck Millipore) with a membrane MWCO of 3 kDa. Filtering of 4 mL medium was proceeded until 100 μL medium was left above the filter. Supernatant and filtrate were recovered and filled up to the initial volume with RPMI-1640.

#### Benzonase treatment

Benzonase was added to conditioned media with a final concentration of 1 U/μL. After that, media were incubated for 30 min at 37 °C.

#### DNase I treatment

DNase I was added to conditioned media with a final concentration of 2,5 U/mL. After that, media were incubated for 30 min at 37 °C. DNase I was inactivated by heating at 75 °C for 10 min.

### Sample preparation for liquid chromatography coupled to mass spectrometry (LC-MS) analysis

Lymph node specimens were weighed into Precellys tubes prefilled with ceramic beads (Stretton Scientific Ltd., Derbyshire, UK). An exact volume of extraction solution (30% acetonitrile, 50% methanol, and 20% water) was added to obtain 60 mg specimen per mL of extraction solution. Samples were lysed using a Precellys 24 homogeniser (Stretton Scientific Ltd., Derbyshire, UK). The suspension was mixed and incubated for 15 minutes at 4 °C in a Thermomixer (Eppendorf, Germany), followed by centrifugation (16,000 g, 15 min at 4°C). The supernatant was collected and transferred into autosampler glass vials, which were stored at −80 °C until further analysis.

Cells (7.5 × 10^4^) were seeded onto 6-well plates and grown for 24h in full growth medium. Medium was then replaced by RPMI without FCS. Control (no cells) and cell-conditioned medium (200 μl) was collected from each well after 24h. The collected media was centrifuged at 4°C for 10 min at 13,200 rpm and 50 μl of the supernatant extracted in 750 μl of cold metabolite extraction solution. Following centrifugation at 4°C for 10 min (13,200 rpm), the supernatant was transferred onto autosampler glass vials and stored as above. Cell-conditioned medium extracts from five independent cell cultures was analysed for each condition, as well as control (non-conditioned) cell culture medium extracts. Samples were randomised in order to avoid bias due to machine drift and processed blindly.

LC-MS analysis was performed using a QExactive mass spectrometer coupled to a Dionex U3000 UHPLC system (Thermo Fisher Scientific). The liquid chromatography system was fitted with a Sequant ZIC-pHILIC column (150 mm × 2.1 mm) and guard column (20 mm × 2.1 mm) from Merck Millipore (Darmstadt, Germany) and temperature maintained at 45 °C. The mobile phase was composed of 20 mM ammonium carbonate and 0.1% ammonium hydroxide in water (solvent A), and acetonitrile (solvent B). The flow rate was set at 200 μL/min with the gradient described previously.^1^ The mass spectrometer was operated in full MS and polarity switching mode. The acquired spectra were analysed using XCalibur Qual Browser and XCalibur Quan Browser software (Thermo Fisher Scientific). Absolute quantification of metabolites in the cell culture medium and LNs was performed by interpolation of the corresponding standard curves obtained from commercially available compounds running with the same batch of samples.

### Statistical Analyses

Statistical analyses were performed using GraphPad Prism 6 software (GraphPad). For comparisons of three or more groups, data were subjected to one-way or two-way ANOVA analysis, followed by post hoc test (Dunnett’s when comparing every mean to a control mean, or Tukey’s multiple comparisons test when comparing every mean to every other mean). When two groups were compared, a two-tailed unpaired Student’s t-test was applied. Data are represented as mean ± SEM, and p ≤ 0.05 was considered significant.

### Cytokine array

Cytokine Arrays were performed according to the R&D System Proteome Profiler Mouse XL Cytokine Array Kit (ARY028, R&D Systems) product’s manuals. Medium from B16.F10 tumour cells was collected containing 2% FBS, 24 h after seeding and centrifuged at 300 xg to remove cell debris. For analysis of tumor lysates, tumors were excised from mice, homogenized in RIPA buffer using a tissue grinder and subsequently spun at 13,000 rpm for 10 min. A BCA (Thermo Scientific) assay of the supernatant was performed to quantify protein content and 1 mg of tumour lysate was used per membrane of the cytokine array. Analysis was done by use of ImageJ with the “Protein Array Analyser” Macro.

### Generation and Analysis of the Computational Model

The computational model was developed in the BioModelAnalyzer (BMA). The BMA allows generation of complex, discrete, executable models using a simple user interface, before models can be interrogated through simulation analysis, stability analysis, and linear temporal logic. Nodes in the model are generally defined as the representing a single protein, gene, or metabolite species, and changes in the discrete value of this node represent alterations in the concentration or activity of that species. In this model most nodes have a range of 0-4, where 2 is defined as normal activity/concentration, 0-1 as low activity/concentration, and 3-4 as high activity/concentration. Where nodes differ from this they have been annotated in the model file, a notable example is the Oxygen Consumption Rate (OCR) node, which ranges from 0-10, from low to high activity respectively. This change was made to more accurately compare values to experiments.

Nodes in the BMA are linked through simple mathematical operations that describe how two nodes relate to each other (called a target function). A target function contains a simple mathematical operation, the result of which describes the value at which a node will tend to. Each step of a simulation of proof analysis, a node may change by a single value (up by one, down by one, or no change), and will tend towards the value of its target function. Nodes are updated in synchrony each step of a simulation. Target functions can encapsulate inputs from multiple sources, and can generate extremely complex regulatory relationships.

The metabolism model was generated through studying of the literature. The core metabolic network is generally well understood, and as such was generated primarily from literature^2^, a summary of the references used in the generation of the model are found in **Table 1**. Metabolic analysis was performed with the use of a special series of nodes. We included a “counter” node, which over the course of a simulation in which a metabolic analysis experiment is run, increases in value from 0 to 100, roughly the number of minutes of a standard metabolic analysis experiment. When the “counter” node reaches particular values, the target functions of other nodes result in their activation, in this way we are able to apply different drug treatments to the model at set intervals, attempting to replicate the temporal properties of the experiments.

---

**Supplementary figure 1.**
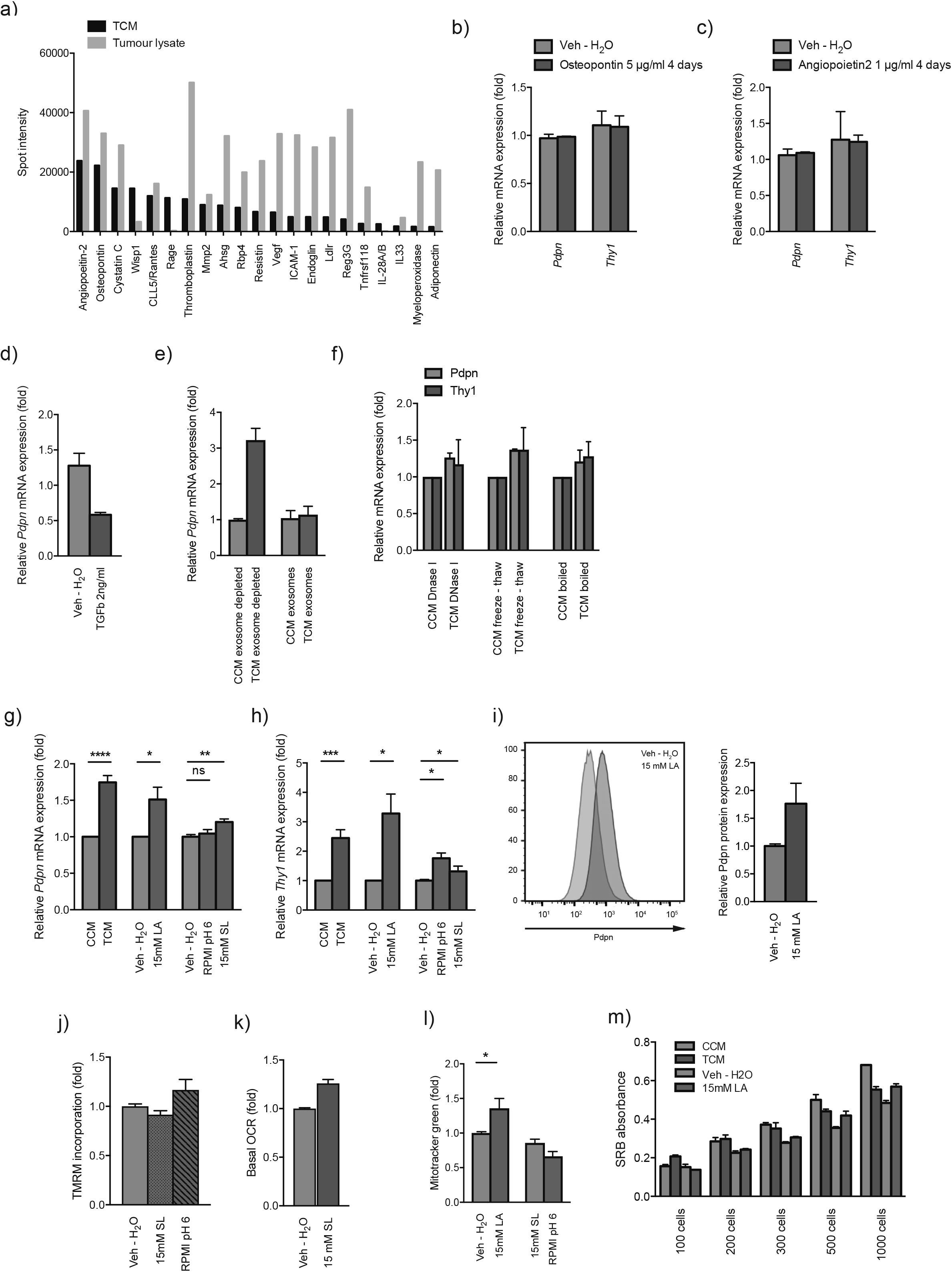
Factor depletion, sodium lactate and low pH medium controls, and proliferation assay. (**A**) Cytokine array of TCM and tumor lysates showing the top 21 detected cytokines as measured spot intensity. (**B** and **C**) Quantitative RT-PCR analysis of *Pdpn* and *Thy1* in *in vitro* cultured FRCs treated for 4 days with 5 μg/ml recombinant murine osteopontin or 1 μg/ml recombinant murine angiopoietin 2. (**D** and **E**) Quantitative RT-PCR analysis of *Pdpn* in *in vitro* cultured FRCs treated for 4 days with 2 ng/ml TGFβ or exosome depleted CCM or TCM or with exosomes isolated from CCM or TCM. (**F**) Quantitative RT-PCR analysis of *Pdpn* and *Thy1* in *in vitro* cultured FRCs treated for 4 days with CCM or TCM treated prior to administration with DNase I, a repeated freeze and thaw cycle or were boiled. (**G** and **H**) Quantitative RT-PCR analysis of *Pdpn* and *Thy1* in FRCs cultured *in vitro* and treated with CCM, TCM, Vehicle (Veh – H_2_O), 15mM lactic acid (LA), RPMI at pH 6 or 15mM sodium lactate (SL) for 4 days. (**I**) Pdpn protein expression in *in vitro* cultured FRCs and treated for 4 days with either vehicle (Veh – H_2_O) or 15mM LA measured by flow cytometry. Representative histogram (left) and relative MFI(right). (**J**) *In vitro* FRCs treated for 4 days with Vehicle (Veh – H_2_O), RPMI at pH 6 or 15mM sodium lactate (SL), stained with TMRM and analyzed by flow cytometry. (**K**) Baseline OCR from vehicle (Veh – H_2_O) and 15mM sodium lactate (SL) treated FRCs. (**L**) *In vitro* FRCs treated for 4 days with vehicle (Veh – H_2_O), 15mM LA, RPMI at pH 6 or 15mM sodium lactate (SL) stained with Mitotracker green and analyzed by flow cytometry. (**M**) SRB proliferation assay of in vitro cultured FRCs treated for 4 days with CCM, TCM, vehicle (Veh – H_2_O) or 15mM LA and seeded in a 96-well plate at different starting cell numbers.

**Supplementary figure 2.**
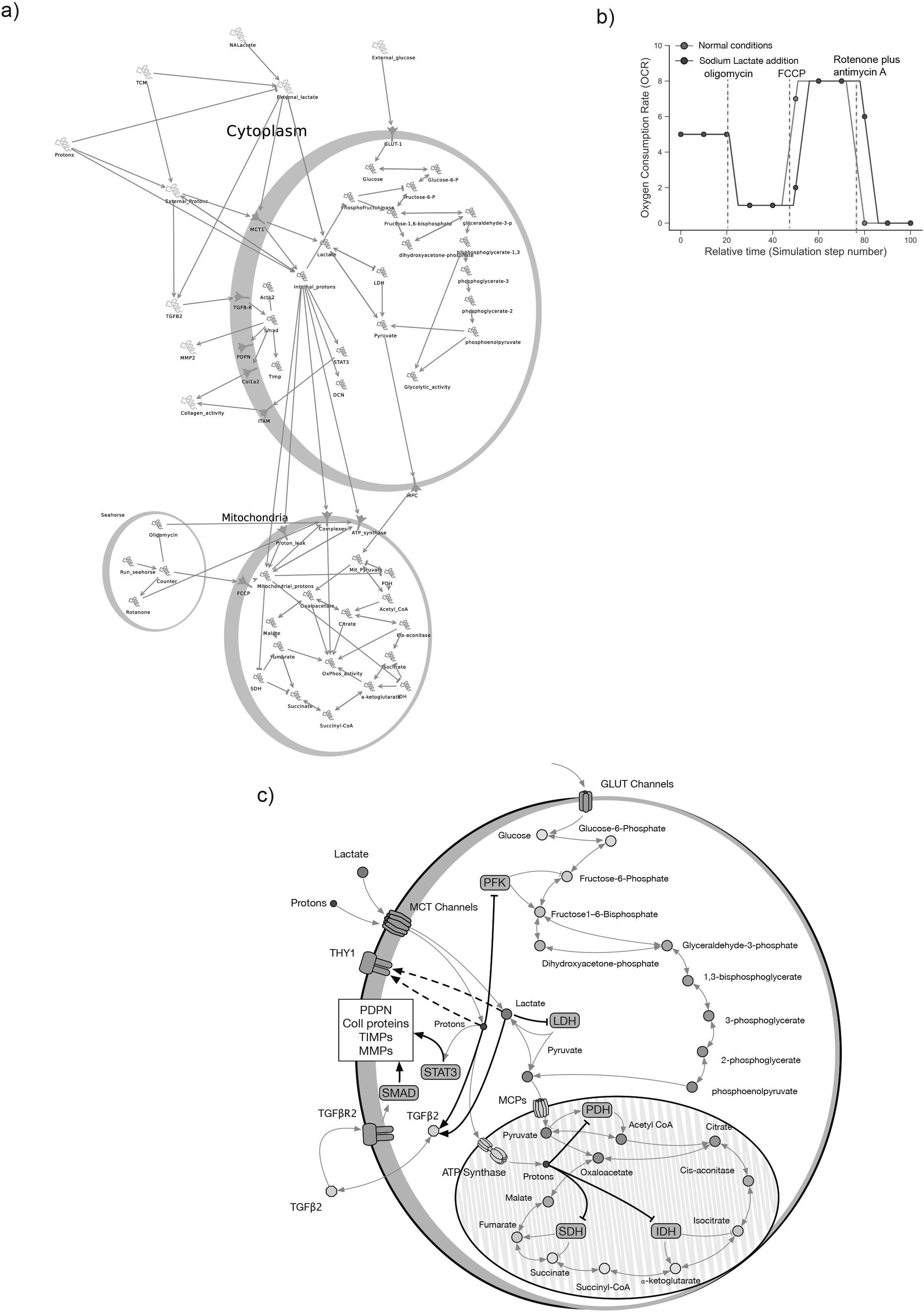
Computational Model of a Metabolic Network. (**A**) Model generated in BioModelAnalyzer (BMA) interface. Shown are “cells” containing cytoplasmic and mitochondrial networks. Model generation is detailed in the methods. (**B**) Metabolic analysis of mitochondrial function in the computational model. Shown is the OCR for mitochondria exposed to sodium lactate compared to control cells. (**C**)Schematic visualizing the findings and model. Lactate and a proton are cotransported into the cell and inhibit not only LDHA and PFK, but also activate Tgfb2, the transcription factors Smad and Stat3, which upregulate Pdpn, collagen synthesis, Timps and Mmps. Protons can also be transported into the mitochondria and the pH shift inhibits the activity of enzymes involved in the TCA cycle. Enzymes are blue boxes, disruptions of normal activity are black arrows.

